# The distribution of epistasis on simple fitness landscapes

**DOI:** 10.1101/490573

**Authors:** Christelle Fraïsse, John J. Welch

## Abstract

Fitness interactions between mutations can influence a population’s evolution in many different ways. While epistatic effects are difficult to measure precisely, important information about the overall distribution is captured by the mean and variance of log fitnesses for individuals carrying different numbers of mutations. We derive predictions for these quantities from simple fitness landscapes, based on models of optimizing selection on quantitative traits. We also explore extensions to the models, including modular pleiotropy, variable effects sizes, mutational bias, and maladaptation of the wild-type. We illustrate our approach by reanalysing a large data set of mutant effects in a yeast snoRNA. Though characterized by some strong epistatic interactions, these data give a good overall fit to the non-epistatic null model, suggesting that epistasis might have little effect on the evolutionary dynamics in this system. We also show how the amount of epistasis depends on both the underlying fitness landscape, and the distribution of mutations, and so it is expected to vary in consistent ways between new mutations, standing variation, and fixed mutations.

## Introduction

Fitness epistasis occurs when allelic variation at one locus affects allelic fitness differences at other loci. Epistatic interactions can be used to uncover functional interactions [1], but for other questions, the most important quantity is the complete distribution of epistatic effects. The shape of this distribution can affect a population’s ability to adapt, its genetic load, the outcomes of hybridization, and the evolution of recombination rate, or investment in sexual reproduction [2–13].

To investigate such questions, most research has focussed on the mean level of epistasis. This can be estimated from the rate at which mean log fitness declines with the number of mutations carried [7,14–17], which is simple to model [2,4,9,18,19]. But variation around this mean can also affect the evolutionary dynamics [6,7,17].

To understand the complete distribution of effects, one approach is to use a simple model of optimizing selection acting on quantitative traits [10,12,20,21]. This approach, which stems from Fisher’s geometric model [22], naturally generates fitness epistasis, even when mutations are additive on the phenotype. Furthermore, the overall level of epistasis can be “tuned” by adjusting the curvature of the fitness function, that is, the rate at which fitness declines with distance from the optimum [10–12,23–28].

Because it generates a rich spectrum of effects with few parameters, Fisher’s geometric model is particularly suitable for fitting to data [24,29–31], including data on fitness epistasis [32–36]. Perhaps most impressively, Martin et al. [32] used the model to successfully predict several properties of the distribution of epistatic effects in the microbes *Escherichia coli* and Vesicular Stomatitis Virus [15,37]. However, these authors did not directly study the effects of varying the curvature of the fitness landscape, and neither did they explore other possible variants of Fisher’s geometric model [25,38–41]. Here, following [32], we study properties of fitness epistasis under Fisher’s geometric model. We extend previous results by examining a wider class of fitness landscapes, and also compare the predictions to a recent, large-scale data set of yeast mutants [1].

## Models and analysis

### Basic notation and a null model without epistasis

Let us denote as ln *w*_*d*_, the log relative fitness of an individual carrying *d* mutations. Across many individuals, the scaled mean and standard deviation of this quantity are

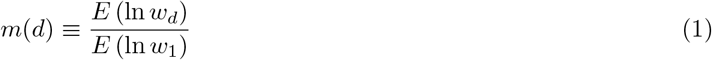

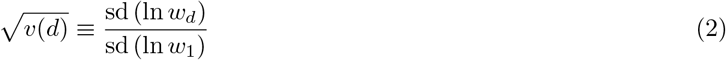

where, by definition, *m*(0) = *v*(0) = 0 and *m*(1) = *v*(1) = 1. These equations use a log scale, because deviations from multiplicativity (i.e. from additivity on a log scale) influence the evolutionary dynamics [7].

We can immediately give results for a null model with no epistatic effects. In this case, mutations will contribute identically to the mean and variance in fitness, regardless of how many other mutations are carried. So a collection of individuals carrying two random mutations are expected to have twice the decline in log fitness, and twice the variance in log fitness, as a collection of individuals carrying one mutation. This implies that

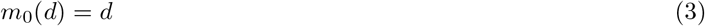

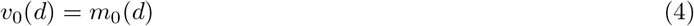

where the subscript 0 indicates the non-epistatic null model. These predictions are illustrated by red lines in Figure 1.

To measure epistasis directly, we could measure the pairwise interaction between two mutations, denoted *a* and *b*:

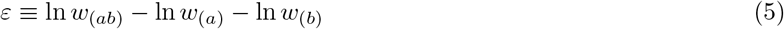

Here, *w*_(*a*)_ denotes the relative fitness of the genome carrying the mutation “*a*”, and so on. Though widely used, *ε* can be difficult to work with. For example, when the same fitness measurements are included multiple times in a data set (e.g., if the same mutation appears in multiple double mutants), then there are complications from pseudoreplication or correlated errors. Furthermore, for a complete picture of epistasis, we would also have to consider higher-order interactions between three or four mutations. For these reasons, in the main text, we will focus on the simpler quantities of eqs. 1–2, and give some equivalent results for *ε* in Appendix 1. The quantities are also closely related. For example, eq. 3 implies that there is no epistasis on average (i.e., that positive effects exactly match negative effects, such that *E* (*ε*) = 0), while eq. 4 states that all epistatic effects are the same, such that Var (*ε*) = 0. Together, then, eqs. 3–4 imply that there is no epistasis at all.

**Figure 1:**
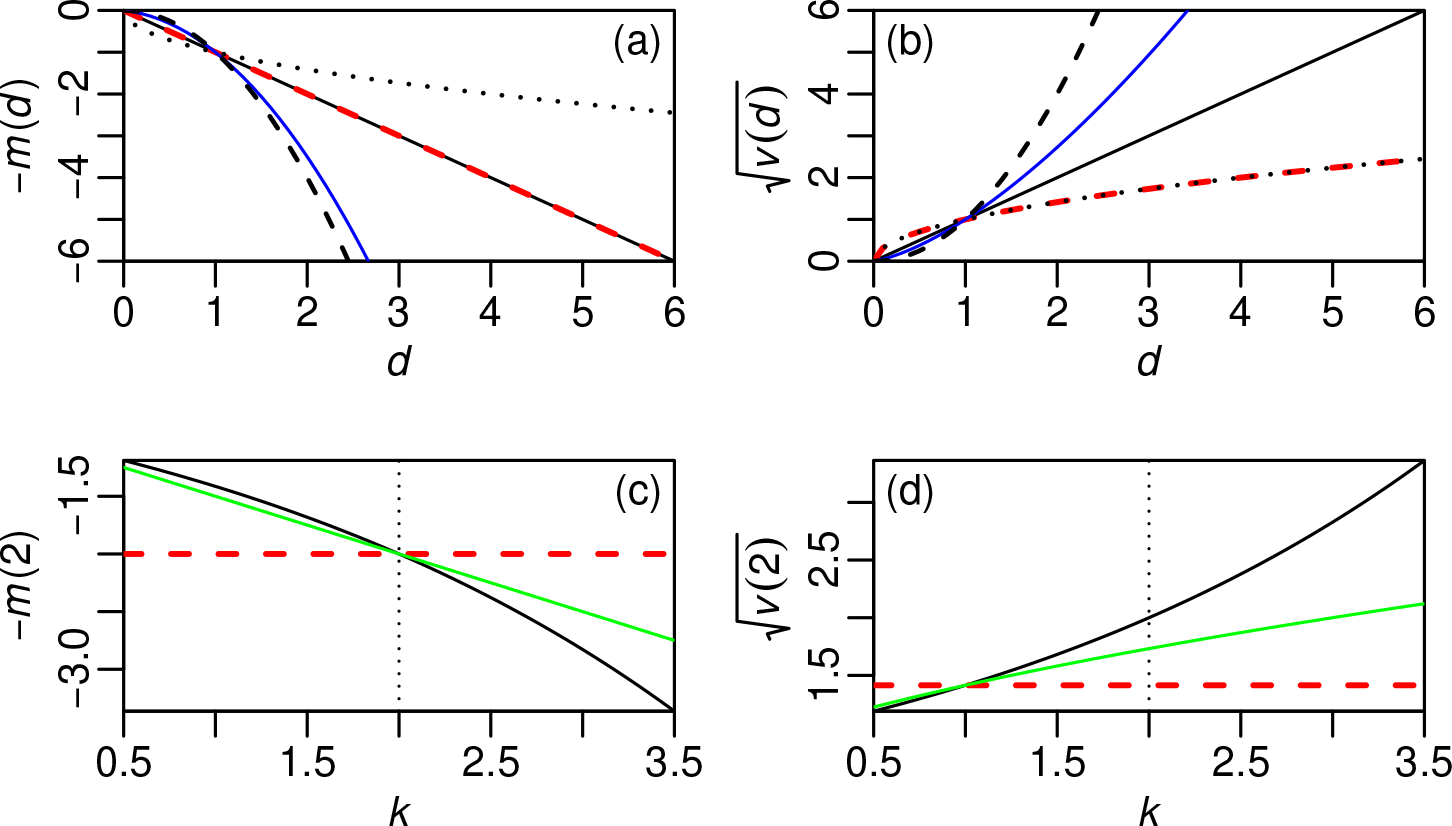
Predictions for mean log fitness (a,c) or the standard deviation in log fitness (b,d). Upper panels show predictions for individuals carrying different numbers of mutations, *d*. Lower panels show results for double mutants (*d* = 2), varying the curvature of the fitness landscape, *k*. Results for the null model, with no epistasis, are shown as red dashed lines. In this case, the mean and variance in log fitness both change linearly with *d* (eqs. 3–4). Results for simple phenotypic models are shown as black lines. The upper panels show results with no epistasis on average (solid lines, *k* = 2), negative epistasis on average (dashed lines, *k* = 4), or positive epistasis on average (dotted lines, *k* = 1). Blue lines show results for a model with strongly biased mutations (*β* = 3, *k* = 2; eqs. 51-52); green lines show results where the mutations on each trait are drawn from a leptokurtic reflected exponential distribution (eqs. 44).

### Additive phenotypic models

We now examine results under Fisher’s geometric model. Here, an individual’s fitness depends on its phenotype, described as an *n*-dimensional vector, **z** = {*z*_1_, *z*_2_,…, *z*_*n*_}, whose components, *z*_*i*_, are the value of each trait. There are several ways in which trait values could combine to yield overall fitness. The simplest is to use their Euclidean distance from the origin, raised to the *k*^th^ power.

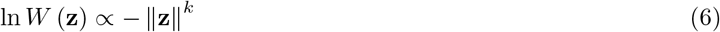

where 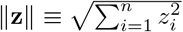 [25,26]. An alternative, which does not assume identical selection on all traits, is

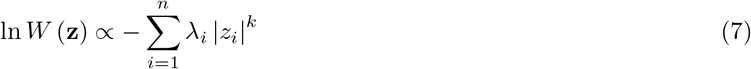

where λ_*i*_ determines the strength of selection on trait *i* [23,24]. These two fitness functions often give similar results (Figures S1-S2), but they are identical only when *k* = 2, and all λ_*i*_ are equal.

The simplest versions of the model make three further assumptions: (1) that the wild-type is phenotypically optimal; (2) that mutations are additive with respect to the phenotype, and (3) that the mutant effects on each trait are drawn, independently, from a standard normal distribution. In this case, the relative fitness of an individual carrying *d* mutations is

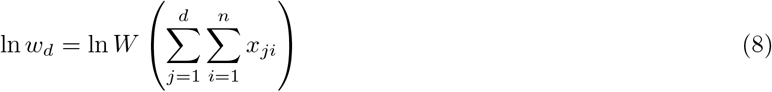

where

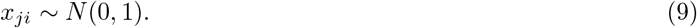

In Appendix 1, we show that, for both fitness functions, these assumptions yield the following results, as illustrated by the black lines in Figure 1:

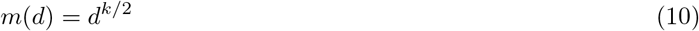

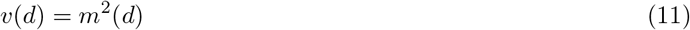

Eqs. 10–11 show how *k* affects the level of fitness epistasis [23,26]. When *k* = 2, we have no epistasis on average, as with eq. 3 (solid black in lines in Fig. 1a-b). Setting *k* > 2 leads to negative epistasis on average (dashed black in lines in Fig. 1a-b), and *k* < 2 leads to positive epistasis on average (dotted black in lines in Fig. 1a-b). Note also, that eq. 11 will never agree with eq. 4, because these simple phenotypic models always generate fitness epistasis.

### Extensions to the phenotypic model

Confronted with data from real quantitative traits [42], many aspects of the model above appear grossly unrealistic. For example, unless the number of traits is very small, the i.i.d. normal model suppresses mutations of overall small effect, and yet there is good reason to think that such mutations are very common [39,43–45].

Furthermore, there is clear evidence that both selection and mutation are correlated among traits [46,47], and that mutations are characterised by highly leptokurtic distributions with stronger concentrations of very small and very large effects; and bias, with a tendency to change traits in a particular direction [48,49]. Furthermore, there is some evidence of appreciable epistasis at the level of phenotype [50,51]; and restricted or modular pleiotropy, where mutations affect only a subset of traits ([39,52]; though see [53]). Finally, there is often evidence of beneficial mutations, which implies that the wild-types are suboptimal. None of this is consistent with eqs. 8–9.

Some of the simplifying assumptions are only apparent. For example, the major effect of correlations can often be transformed away, by redefining the axes, and considering a smaller “effective number of traits” [21,29,46]. Nonetheless, other assumptions are certainly restrictive. In Appendix 1, we explore several extensions to the model, focussing on assumptions that can be relaxed in a general way, and building on results from previous studies [29,32,38,39,41,44,46]. In particular, we consider variable distributions of effect sizes, restricted pleiotropy, mutational bias, and suboptimal wild-types. Despite their heterogeneity, most of these extensions act to reduce mean levels of epistasis. With modular pleiotropy, this is because mutations affecting different traits will interact less; with high kurtosis, it is because epistasis is reduced when any of the mutations is very small in magnitude; finally, parental maladaptation reduces “overshoots” of the optimum, which cause sign epistasis [27]. In all cases, the predicted *m*(*d*) is intermediate between predictions from the simplest phenotypic models (eq. 10) and the null model (eq. 3). This is illustrated by the green lines in Figure 1c, which show results with a leptokurtic distribution of effects on each trait. Only one of the modifications has a qualitatively different effect. When mutations are biased, their tendency to modify traits in a consistent direction makes epistasis more negative. To illustrate this, let us assume that mutational effects have a non-zero mean, *β_i_*, such that, *x_ij_* ~ *N*(*β_i_*, 1). When the bias is large, we find that

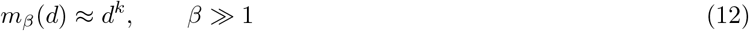

where 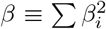 (see Appendix 1 for details). The decline of the mean fitness is now more rapid than in a model without bias (compare eqs. 10 and 12), and this is illustrated by the blue lines in Figure 1a, which show the effects of bias when *k* = 2.

For the variance in log fitness, the effects are even more consistent. For all of the extensions, we find a reduction, compared to simplest phenotypic model, such that

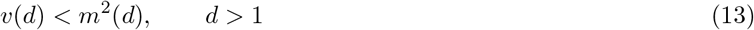

and when *k* ≥ 2, results for the null model act as lower bound, such that *v*(*d*) ≥ *m*(*d*). This is illustrated by the green and blue lines in Figure 1b and d.

To summarize, modifying the phenotypic model, to reflect data from real quantitative traits, has two main effects. First, it erases information about the true curvature of the fitness landscape, so that the form of *m*(*d*) cannot easily be used to estimate *k*. Second, it reduces the variance in log fitness, below *m*^2^(*d*).

### Reanalysis of data from a yeast snoRNA

To illustrate the approach above, we now reanalyse data from Puchta et al. [1], who used saturation mutagenesis of the U3 snoRNA in *Saccharomyces cerevisae* (see Appendix 2 for full details). Figure 2a confirms that pairwise epistatic interactions are present in these data [1]. Nevertheless, Figure 2c-d show that, considered as a whole, the data give a very good fit to the non-epistatic null model (eqs. 3–4).

**Figure 2:**
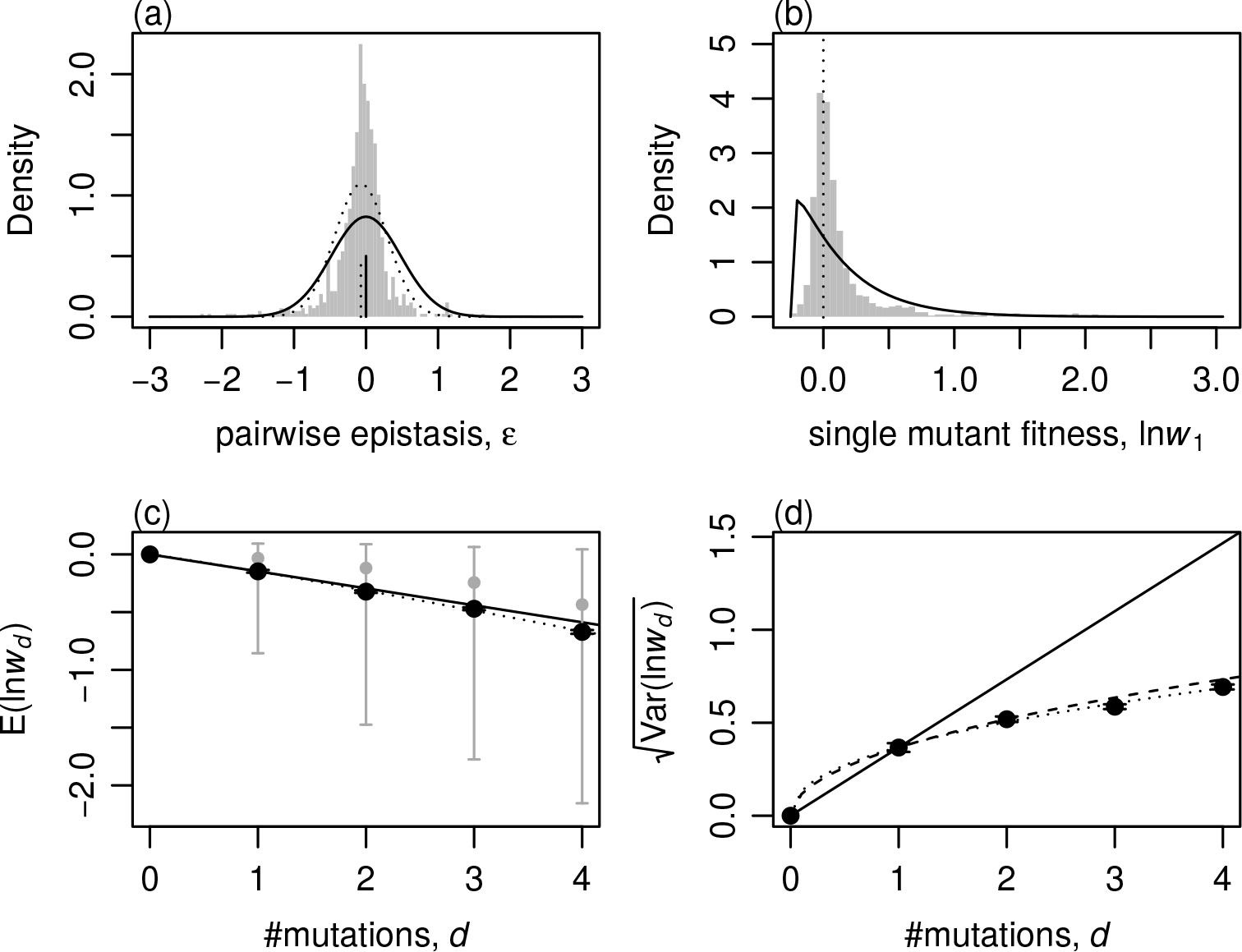
Reanalysis of mutations in *Saccharomyces cerevisiae* U3 snoRNA [1]. (a) shows the distribution of pairwise epistatic effects (eq. 5), compared to the predictions of the simplest phenotypic model: *ε* ~ *N*(0, 2Var (ln *w*_1_)) (black line [32]; Appendix 1), and a normal distribution with matching mean and variance (dotted line). (b) shows the distribution of single mutant log fitnesses, and the best-fit shifted gamma distribution, as predicted by the simplest phenotypic models [29]. (c) shows the mean of the log fitnesses of individuals carrying *d* mutations (black points with standard error bars); the median and 90% quantiles (grey points and bars); the analytical prediction, which applies to both the null model and the phenotypic model with *k* = 2 (black line; eqs. 3 and 10); and the best-fit regression for ln *m*(*d*) ~ ln *d* (dotted line, which has a slope implying *k̂* = 2.16). (d) shows the standard deviation in the log fitnesses of individuals carrying *d* mutations (black points with standard error bars); the median and 90% quantiles (grey points and bars); analytical predictions from the null model, eq. 3 (dashed line), or the phenotypic model with *k* = 2, eq. 10 (solid line); and the best-fit regression of ln *v*(*d*) ~ ln *d* (dotted line, which has slope 0.89).

Some of this apparent discrepancy can be attributed to the greater robustness of our statistics to measurement error. For example, we show in supplementary Figures S4 and S5, that the inferred variance in epistatic effects decreases with the amount of replication, while patterns in *m*(*d*) and *v*(*d*) are little changed. Furthermore, some reduction in epistasis, relative to simple phenotypic model, could have been predicted from other aspects of the data. For example, the distribution of single-mutant fitnesses (Figure 2b), shows that the distribution is highly leptokurtic, and indicates the presence of beneficial mutations (346/965 mutations increase growth rate). Nevertheless, kurtosis and wild-type maladaptation both need to be extreme for predictions to converge to the null model (see Appendix 1). Furthermore, the hypothesis of modularity, whereby mutations each affect different sets of traits, seems inherently implausible for these data, where all mutations affect sites in the same snoRNA. As such, we conclude that the phenotypic models - even in modified form - overestimate the true amount of fitness epistasis in these data. This implies that the simplest population genetic models, which ignore epistasis altogether, might be sufficient to understand the evolutionary dynamics in this system, despite the clear presence of some fitness interactions [1].

## Discussion

We have used simple summary statistics to describe levels of fitness epistasis. These statistics are relevant to evolutionary questions [7], and are less sensitive to measurement error than are estimates of individual epistatic effects.

We then developed analytical predictions for these statistics under simple models of selection on quantitative traits. The simplest such model assumes that mutant effects on each trait are i.i.d. normal, and this seems unrealistic [39,42–44]. Nevertheless, this model, with *k* = 2, has been shown to give a good fit to fitness data from *E. coli* and VSV [15,32,37]. Our results go further, and show that only this simple model would have fit those data; increasing the realism of the quantitative traits (e.g., by introducing leptokurtic effects, or restricting pleiotropy), would have underpredicted the amount of epistasis. This finding has two implications. First, the success of the “unrealistic” model, reinforces the argument of [21], that the traits in Fisher’s geometric model, when considered as a fitness landscape, should not be equated with standard quantitative traits. Second, multiple authors have shown that models with no epistasis on average (i.e., with *k* = 2), are vulnerable to invasion by modifiers [26,54,55], and so the good fit of *k* = 2 implies that global modifiers of fitness epistasis do not arise in these systems.

Of course, there is no reason to assume that identical patterns of epistasis will characterise all data sets [56,57], and we have offered two further reasons to doubt this. First, empirically, we have shown that the data of [1] give a good overall fit to a non-epistatic null model, despite the clear presence of some fitness interactions (Figure 2). Second, theoretically, we have shown how the observed level of epistasis will depend on both the underlying fitness landscape, and the distribution of mutation effects. For example, a landscape with a high level of curvature (i.e., *k* > 2), might still generate a linear decline in mean log fitness (such that *m*(*d*) ≈ *d*) if the distribution of mutant effect sizes is highly leptokurtic, but this effect should be evident in the reduced levels of variance (such that *v*(*d*) < *m*^2^(*d*) for *d* > 1). If the fixation process acts to restrict the distribution towards mutations of medium size [38], the levels of observed epistasis should therefore increase steadily for new mutations, standing variation, and differences that are fixed between populations.

## Supporting information

## Acknowledgements

We are very grateful to Grzegorz Kudla, Anna Puchta, Elena Kuzmin, Santiago Elena and Rafael Sanjuan for providing their data, and helpful clarifications. We are also grateful to Denis Roze, Nicolas Bierne, Chris Illingworth, and Fyodor Kondrashov for useful discussions.

## Author contributions

Both authors designed the study, analysed the data and wrote the manuscript. JW carried out the modelling.

