## Supplementary material for "The distribution of epistasis on simple fitness landscapes"

### Appendix 1: Derivations

In this Appendix, we derive the key results in the main text, and justify claims about the extensions to the simplest phenotypic model. We will also present results for direct measures of pairwise epistasis (eq. 5).

#### 1. The distribution of pairwise epistatic effects:

Martin et al. [1] examined the scaled moments of the distribution of pairwise epistatic effects (eq. 5). These moments are closely related to the scaled moments of the genotypic fitness values, $m(d)$ and $v(d)$, that we use in the main text (eqs. 1-2). To see this, let us consider individuals with a wild-type phenotype $\mathbf{z}$. The relative fitness of an individual carrying a single mutation, with phenotypic effects $\mathbf{x}$, is

$$\ln w_1 \equiv \ln W(\mathbf{z}+\mathbf{x}) - \ln W(\mathbf{z}) \tag{14}$$

This is closely related to the selection coefficient of the mutation, $s$, because when $s$ is small in magnitude, $s \approx \ln(1+s) = \ln w_1$. This is why the quantity shown in eq. 14 is denoted as $s$ by Martin et al. [1]. From eq. 5, the pairwise epistatic effect for two mutations, $a$ and $b$, is then

$$\varepsilon = \ln W(\mathbf{z}+\mathbf{x}_a+\mathbf{x}_b) - \ln W(\mathbf{z}+\mathbf{x}_a) - \ln W(\mathbf{z}+\mathbf{x}_b) + \ln W(\mathbf{z}) \tag{15}$$

[1,2]. We can now use eqs. 1-2 to write the mean and variance of epistatic effects, scaled by the same quantities for single mutations:

$$\frac{E(\varepsilon)}{E(\ln w_1)} = m(2) - 2 \tag{16}$$

$$\frac{\mathrm{Var}(\varepsilon)}{\mathrm{Var}(\ln w_1)} = v(2) + 2 - 4\sqrt{v(2)}\,r_{12} \tag{17}$$

Here, we have defined

$$r_{12} \equiv \mathrm{Cor}\left(\ln W(\mathbf{x}_a+\mathbf{x}_b+\mathbf{z}), \ln W(\mathbf{x}_a+\mathbf{z})\right) \tag{18}$$

as the correlation coefficient between the fitnesses of genotypes carrying a single mutation alone, and in combination with a second mutation. Under the null model, with no epistasis, the assumption that $\mathrm{Var}(\varepsilon) = 0$, combined with eq. 4, implies that $r_{12} = \sqrt{1/2} \approx 0.707$, and so this value can be compared to results from other models below.

#### 2. Results for the simplest phenotypic model:

Let us first consider results for the simplest model, when the wild-type is phenotypically optimal ($\mathbf{z} = \mathbf{0}$), and the effects of each mutation on each trait are drawn from independent standard normal distributions (eq. 9).

If we use the fitness function of eq. 6 [2,3], which assumes equal selection on all $n$ traits, then the quantities we require for eqs. 1-2 are simply moments of the Chi-squared distribution, with $n$ degrees of freedom:

$$E\left(\ln w_d\right) = (2d)^{k/2}\frac{\Gamma\left(\frac{k+n}{2}\right)}{\Gamma\left(\frac{n}{2}\right)} \tag{19}$$

$$\mathrm{Var}\left(\ln w_d\right) = (2d)^{k}\left(\frac{\Gamma\left(\frac{2k+n}{2}\right)}{\Gamma\left(\frac{n}{2}\right)} - \frac{\Gamma^2\left(\frac{k+n}{2}\right)}{\Gamma^2\left(\frac{n}{2}\right)}\right) \tag{20}$$

The results of eqs. 10-11 follow directly.

If we allow for variation in the strength of selection between traits, and use the fitness function of eq. 7 [4], then the key quantities are now the moments of a folded normal distribution (i.e., the absolute value of a normally-distributed random variable).

$$E\left(\ln w_d\right) = (2d)^{k/2}\frac{\Gamma\left(\frac{k+1}{2}\right)}{\sqrt{\pi}}\sum_{i=1}^{n}\lambda_i \tag{21}$$

$$\mathrm{Var}\left(\ln w_d\right) = (2d)^{k}\left(\frac{\Gamma\left(\frac{2k+1}{2}\right)}{\sqrt{\pi}} - \frac{\Gamma^2\left(\frac{k+1}{2}\right)}{\pi}\right)\sum_{i=1}^{n}\lambda_i^2 \tag{22}$$

and again, eqs. 10-11 follow directly. Figure S1a confirms, with simulations, that the two fitness functions give identical results.

##### 2.1 Pairwise epistatic effects

To calculate the variance in pairwise epistatic effects (eq. 17), we also require the correlation coefficient of eq. 18. For the fitness function of eq. 7, this is maximized at $k = 2$, where it takes the value:

$$r_{12} = \mathrm{Cor}\left(\left|x_{ia} + x_{ib}\right|^2, \left|x_{ia}\right|^2\right) = \frac{1}{2}, \quad k = 2 \tag{23}$$

and so the correlation between single- and double-mutant fitnesses is always lower than under the null model. The same value holds approximately for other values of $k$, and for the alternative fitness function of eq. 6. As such, we have the results

$$-\frac{E\left(\varepsilon\right)}{E\left(\ln w_1\right)} = 2 - 2^{k/2} \tag{24}$$

$$= 0, \quad k = 2 \tag{25}$$

$$\frac{\mathrm{Var}\left(\varepsilon\right)}{\mathrm{Var}\left(\ln w_1\right)} \approx 2(1 + 2^{k-1} - 2^{k/2}) \tag{26}$$

$$= 2, \quad k = 2 \tag{27}$$

These results are compared to simulation in Fig. S2. When $k = 2$, eqs. 25 and 27 reproduce the results of Martin et al. [1], while increasing $k$ above this value makes expected levels of epistasis more negative ($E\left(\varepsilon\right) < 0$), and increases the variance in epistatic effects ($\mathrm{Var}\left(\varepsilon\right) > 2\mathrm{Var}\left(\ln w_1\right)$).

The complete distribution of epistasis is also derivable for $k = 2$, since we have

$$\varepsilon \propto \sum_i^n \lambda_i \xi_i, \qquad k=2 \tag{28}$$

where $\xi_i \equiv x_{ai} x_{bi}$, and this has the pdf

$$\text{pdf}(\xi) = \int_0^\infty \frac{\cos(|\xi| t)}{\pi\sqrt{t^2+1}} dt$$

which has a vanishing mean and unit variance. Central limit behaviour means that $\varepsilon$ will be approximately normally distributed when $k = 2$ [1]. As shown in Fig. S2, the mode of the distribution remains close to zero for all $k$ values, meaning that variation in the curvature of the fitness landscape acts to skew the distribution of epistatic effects.

#### 3. Extensions to the simplest phenotypic model

In this section, we consider various extensions to the simplest phenotypic model. These analyses support eqs. 12 and 13 and statements in the main text.

##### 3.1. Modular pleiotropy and variable effects sizes:

The first set of extensions are most easily made with the isotropic fitness function of eq. 6.

Let us first consider the effects of restricting pleiotropy. Instead of assuming that each mutation affects all $n$ traits, we now assume that pleiotropy is modular ([5]; see also [6,7]), such that each new mutation affects a distinct "module" containing $n'$ traits, which are under selection independently of other modules. To treat this case, consider the total length of the phenotypic effect for a double mutant. This can be written as:

$$\|\mathbf{x}_a + \mathbf{x}_b\| = \sqrt{\sum_i^n (x_{ai} + x_{bi})^2} = \sqrt{\|\mathbf{x}_a\|^2 + \|\mathbf{x}_b\|^2 + 2\,\|\mathbf{x}_a\|\,\|\mathbf{x}_b\| \cos(\theta)} \tag{29}$$

where $\theta$ is the angle in radians between the two mutational vectors, in the $n$-dimensional trait space [5]. If the mutations affect different modules, then their individual vectors will be orthogonal, such that $\cos(\theta) = 0$. Since the sum of Chi-squared random variables is also Chi-squared distributed, we require the moments of a Chi-squared distribution, with $dn'$ degrees of freedom:

$$E\left(\ln w_d\right) = 2^{k/2} \frac{\Gamma\left(\frac{k+dn'}{2}\right)}{\Gamma\left(\frac{dn'}{2}\right)}, \tag{30}$$

$$= dn', \qquad k=2$$

$$\text{Var}\left(\ln w_d\right) = 2^k \left( \frac{\Gamma\left(\frac{2k+dn'}{2}\right)}{\Gamma\left(\frac{dn'}{2}\right)} - \frac{\Gamma^2\left(\frac{k+dn'}{2}\right)}{\Gamma^2\left(\frac{dn'}{2}\right)} \right) \tag{31}$$

$$= 2dn', \qquad k=2$$

When $k = 2$, these results immediately reproduce the null model (eqs. 3-4). We also have the approximation

$$\frac{v(d)}{m^2(d)} = \frac{\Gamma(dn'/2+k)\,\Gamma(dn'/2)\,\Gamma^{-2}(dn'/2+k/2)-1}{\Gamma(n'/2+k)\,\Gamma(n'/2)\,\Gamma^{-2}(n'/2+k/2)-1} \approx \frac{1}{d} \tag{32}$$

and, using the Beta function, we have the limits:

$$m(d) = \frac{B\left(\frac{n'}{2}, \frac{k}{2}\right)}{B\left(\frac{dn'}{2}, \frac{k}{2}\right)} \tag{33}$$

$$= d^{k/2}, \quad n' \to \infty \tag{34}$$

$$= d, \quad n' \to 0 \tag{35}$$

More complete models would have to specify the probability that a pair of mutations appears in the same module, and also consider modules of different sizes. However, the results above are sufficient to show that $m(d)$ will be intermediate between the simple phenotypic model eq. 10, and the null model eq. 3, and that eq. 13 will hold. Simulations with intermediate values of $n'$ are shown in Figure S1b, and confirm these claims.

The effects of modular pleiotropy can also be replicated in a model with universal pleiotropy, if we allow for mutations of very different sizes. This is equivalent to assuming a highly leptokurtic distribution of effects on the overall size of mutations, and thereby on each trait. This is easiest to demonstrate by considering pairwise epistatic effects, when $k = 2$. In this case, we have

$$\varepsilon = 2\,\|\mathbf{x}_a\|\,\|\mathbf{x}_b\|\cos(\theta), \qquad k = 2 \tag{36}$$

As shown by Fisher [8], when the number of traits, $n$, is not very small, then an unbiased distribution of mutation directions leads to $2\cos(\theta) \sim N(0, 4/n)$ [9,10]. As such, we have

$$E(\varepsilon) = 0, \qquad k = 2$$

$$\mathrm{Var}(\varepsilon) \approx \frac{4}{n} E\left((\|\mathbf{x}_a\|\,\|\mathbf{x}_b\|)^2\right), \qquad k = 2 \tag{37}$$

If we follow Lourenço et al. [7] and draw the squared mutation magnitudes from a Chi-squared distribution with $n'$ degrees of freedom, then it follows that $\mathrm{Var}(\varepsilon) = \frac{4}{n} n'^2$. The excess kurtosis of the chi-squared distribution is $12/n^{'}$ and so decreasing $n'$ increases the kurtosis, and decreases the variance in epistatic effects. Simulation results, shown in Figure S1c and Figure S2c-d, show that the same general pattern holds for other values of $k$, and for other leptokurtic distributions of mutation sizes.

##### 3.2. Varying the distribution of effects on each trait

In the previous section, we used a "top-down" approach to mutation, in which the vector size and direction were independently calculated [11]. The alternative, "bottom-up" approach is to directly specify the distribution of effects on individual traits. This is simplest with the fitness function of eq. 7, where analytical results can be obtained for double mutants, with $d = 2$.

Because the distribution of mutations on quantitative traits is often leptokurtic, let us first consider results when mutational effects are drawn from a reflected exponential distribution, with parameter $\mu$. In this case, the absolute effect on a single trait, $|x|$, is exponentially distributed, such that

$$E[|x|^k] = k!\mu^k \tag{38}$$

$$\mathrm{Var}\left(|x|^k\right) = \mu^{2k}\left((2k)! - (k!)^2\right) \tag{39}$$

For quantities involving two mutations ($d = 2$), if their effects have the same sign, then we have an Erlang distribution:

$$E\left(|x_a + x_b|^k \, |x_a x_b > 0\right) = \mu^k \frac{\Gamma(2+k)}{\Gamma(2)} = \mu^k (k+1)! \tag{40}$$

If they have different signs, we have a difference in exponentials, whose pdf is

$$f(\delta) = \frac{2}{\mu^2} e^{-|\delta|/\mu} \tag{41}$$

$$E\left(|x_a + x_b|^k \, |x_a x_b < 0\right) = k!\mu^k \tag{42}$$

The signs differ with 50% probability, and so, combining these results, we have

$$\begin{aligned} E[|x_a + x_b|^k] &= \mu^k \left(\frac{(k+1)! + k!}{2}\right) = \mu^k k! \left(\frac{k+2}{2}\right) \\ \mathrm{Var}[|x_a + x_b|^k] &= \mu^{2k}\left[(2k)!\,(k+1) - \left(k!\left(\frac{k+2}{2}\right)\right)^2\right] \end{aligned} \tag{43}$$

and so, we find:

$$\begin{aligned} m(2) &= 1 + \frac{k}{2} \\ v(2) &= \frac{(2k)!\,(k+1) - (k!)^2\left(\frac{k+2}{2}\right)^2}{(2k)! - (k!)^2} \approx 1 + k \end{aligned} \tag{44}$$

where the approximate expression for $v(2)$ uses Stirling's approximation: $k! \approx \sqrt{2n\pi}\left(\frac{n}{e}\right)^n$, such that $(2k)!/(k!)^2 \approx 2^{2k}/\sqrt{\pi k}$. The results are supported by simulations shown in Figure S1d. The important point is that the introduction of kurtosis reduces the curvature in $m(d)$, taking it closer to the null model, while, for the variance, $v(d)$ we have $m^2(2)/v(2) \approx 1 + k^2/(4(1+k))$, such that eq. 13 holds.

For completeness, and to highlight the role of kurtosis, let us now assume a platykurtic distribution of effects, such that the effect on each trait is assumed to be uniformly distributed with mean zero: $x_i \sim U(-u/2, u/2)$. The key quantities can now be found by direct integration for $d = 1$ and $d = 2$.

$$
\begin{aligned}
E\left(\ln w_d\right) &= \frac{(u/2)^k}{k+1}\sum_{i=1}^{n}\lambda_i, \qquad d=1 \\
&= \frac{u^k}{\binom{k+2}{2}}\sum_{i=1}^{n}\lambda_i, \qquad d=2
\end{aligned}
$$

$$
\begin{aligned}
\mathrm{Var}\left(\ln w_d\right) &= (u/2)^{2k}\left((2k+1)^{-1}-(k+1)^{-2}\right), \qquad d=1 \\
&= u^{2k}\left(\binom{2k+2}{2}^{-1}-\binom{k+2}{2}^{-2}\right), \qquad d=2 \qquad (45)
\end{aligned}
$$

and so

$$
m(2) = \frac{2^{k+1}}{k+2}
$$

$$
v(2) = \frac{2^{2k}(k+5)}{(k+2)^2} \qquad (46)
$$

Simulations of this model are shown in Figure S1e. The results show that reducing the kurtosis of the mutational effects acts to increase the effects of epistasis on the mean fitness (i.e., exaggerating the effects of $k$ on $m(d)$), and also increases the variance, such that $v(2) > m^2(2)$.

##### 3.3. Biased mutations, and suboptimal wild-type

In this section, we allow for bias in the effects of mutations (i.e. a non-vanishing mean effect), and relax the assumption that the wild-type genotype, carrying no mutations, is phenotypically optimal. In both cases, this is easiest if we assume the isotropic fitness function of eq. 6.

For bias, we assume that the effects of the $j^{\mathrm{th}}$ mutation on the $i^{\mathrm{th}}$ trait is distributed as

$$
x_{ij} \sim N(\beta_i, 1) \qquad (47)
$$

For suboptimality, we denote as $z_i$, the deviation from the optimum for the $i^{th}$ trait in the wild-type. In this case, the key quantities follow from the moments of a non-central Chi-squared distribution with parameter $\alpha \equiv \sum_i^n \left(z_i + d\beta_i\right)^2$. These moments are given by

$$
\begin{aligned}
E\left(\left(\ln w_d + \ln w_0\right)^P\right) &= (2d)^{Pk/2} e^{-\alpha/(2d)} \frac{\Gamma\left(\frac{Pk+n}{2}\right)}{\Gamma\left(\frac{n}{2}\right)} K\left(\frac{Pk+n}{2}, \frac{n}{2}, \frac{\alpha}{2d}\right) \qquad (48) \\
&= dn + \alpha, \qquad Pk = 2 \\
&= 2d(dn+2\alpha) + (dn+\alpha)^2, \qquad Pk = 4
\end{aligned}
$$

where

$$K(a,b,z) \equiv \sum_{i=0}^{\infty} \frac{\binom{a}{i}}{\binom{b}{i}} \frac{z^i}{i!}$$

is Kummer's confluent hypergeometric function [12]. Simple results now follow for $k = 2$. For general $k$, well defined limits [12], allow us to derive results where maladaptation, or bias, are large.

First, let us consider the case where mutations are unbiased ($\beta_i = 0$), but the wild-type is suboptimal. If we define $\xi = \sum z_i^2$, and note that $\ln w_0 = -\xi^{k/2}$, then we find

$$m_\xi(d) = d, \quad k = 2,$$

$$= d, \quad \xi \to \infty \tag{49}$$

$$v_\xi(d) = d^2 \frac{1 + 2\xi/d}{1 + 2\xi}, \qquad k = 2$$

$$= d = \frac{m_\xi^2(d)}{d}, \quad \xi \to \infty \tag{50}$$

These results show that the non-epistatic null model is approached as the wild-type becomes very maladapted [13].

Results with bias, but an optimal wildtype ($z_i = 0$), follow in the same way. If we define $\beta \equiv \sum \beta_i^2$, then we find:

$$m_\beta(d) = d \frac{1 + d\beta}{1 + \beta}, \quad k = 2$$

$$= d^k, \quad \beta \to \infty \tag{51}$$

$$v_\beta(d) = d^2 \frac{1 + 2d\beta}{1 + 2\beta}, \quad k = 2$$

$$= d^{2k-1} = \frac{m_\beta^2(d)}{d}, \quad \beta \to \infty \tag{52}$$

Note that eqs. 50 and 52, are equivalent to eq. 32, showing that extreme levels of maladaptation, modularity and bias have identical effects on the variance. Simulation results with mutational bias are shown in Figure S1f.

### Appendix 2: Details of data reanalysis

We searched the literature for data sets combining replicated measures of fitness for multiple mutations, chosen without regard for their fitness consequences. We rejected many excellent data sets where the trait measured was not a plausible proxy for fitness [14,15], or which contained no genotypes carrying four or more mutations [16,17], or mutations that were known in advance to be beneficial [18,19], or were otherwise biased [17], or which contained clear edge effects that could not be easily corrected [17,20]. Moreover, we did not consider mutation accumulation lines, where the number of mutations must be estimated, so that estimates can be confounded by changes in mutation rate [21].

For the data set of Puchta et al. [22], a 333-nucleotide long U3 snoRNA gene in *S. cerevisae* was the target of a saturation mutagenesis experiment. The wild-type used was a D343 yeast strain in which the U3 gene was transformed to allow the yeast to survive on a selected environment containing glucose (otherwise U3 is down-regulated and growth arrested). Libraries of U3 mutated strains were constructed using "doped oligonucleotides" that randomly mutated any possible site between position 7 to 333 of the gene (327/333 sites, with ~1% mutation rate per position). All possible point mutations of the U3 gene were represented in the libraries, which contained single-nucleotide polymorphisms (SNPs) and short deletions and insertions (indels). To measure fitness, competition experiments were performed in an environment containing glucose. Following Puchta et al. [22], our main text reports results from the "env. 1" condition, which was kept at 30°C.

Due to the mutagenesis procedure, many mutation combinations were present multiple times, and where this was the case, we averaged over the log fitness estimates. Figure S3a compares the mean and standard deviation of the log fitness estimates for replicated strains. The plot shows a clear trend for heteroscedasticity, with larger fitness effects associated with greater measurement uncertainty (i.e., higher environmental variance). Such heteroscedasticity should increase $v(d)$ above its true value, militating against a fit to the null model, and therefore making our conclusions conservative.

The data set of Puchta et al. [22] also includes additional replication, because fitness estimation was repeated in a second environment at 37°C ("env. 2"), and a third environment, also at 30°C ("env. 3"). As shown in Figure S4a-b, results for the two identical environments were highly correlated. Considering these replicate experiments, clarifies a disadvantage of using direct estimates of pairwise epistasis, eq. 5, because the estimates of this quantity, as shown in Figure S4c, are much less precisely replicated than the estimates of single- or double-mutant effects (Figure S4a-b). Furthermore, the estimated variance in epistatic effects, which was the subject of predictions by Martin et al. [1], is highly sensitive to the amount of replication. This is shown in Figure S4d. By contrast, as shown in Figure S5, the patterns evident in the moments of $\ln w_d$, are relatively robust between the three experiments (Figure S5a), and even more so, when multiple experiments are treated as replicates (Figure S5b). This remains true when we consider only Single Nucleotide Polymorphism mutations (i.e., excluding small insertions and deletions), of the kind that are used in the calculation of pairwise epistasis measures (Figure S5c). As is clear from Figure S5a,c, the experiment "env. 1", which we report in the main text, shows the largest deviations from expectations under the null model, again making our conclusions conservative.

A final consequence of the saturation mutagenesis procedure was that around half of the strains contained more than $d = 4$ mutations, and some contained as many as $d = 57$. We did not reanalyze these highly mutated strains, due to experimental difficulties in measuring very low fitness values. In particular, Puchta et al. [22] truncated their fitness measurements at $\ln w = -3$. This leads to edge effects that are clearly visible in Figure S3b (where log fitness values were averaged across all three replicate experiments). The edge effects are also visible in Figure S6, where we replicate Figure 1a-b, but retaining strains carrying up to $d = 12$ mutations (thereby including ~93% of the data set). These effects explain our conservative choice to restrict the reanalysis to strains carrying $d \leq 4$ mutations in the main text.

### Supplementary Figures

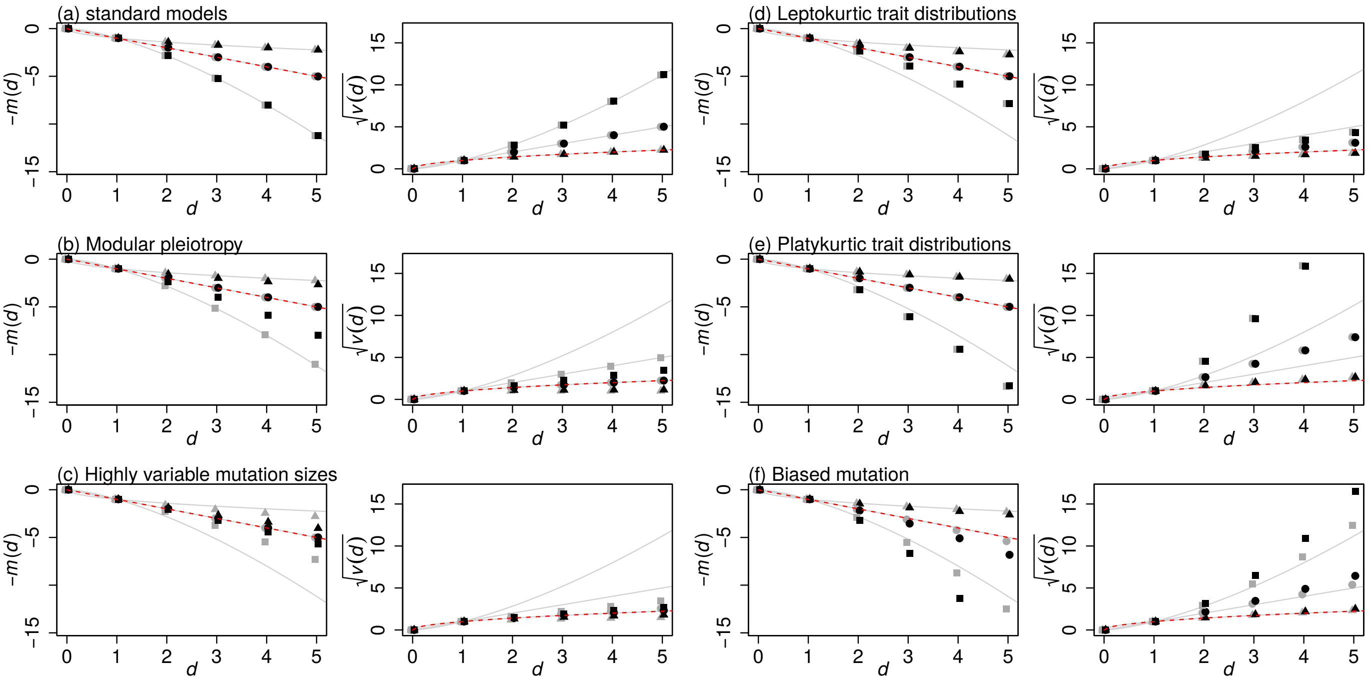


**Figure S1**: Properties of fitness epistasis between mutations under simple phenotypic models, based on Fisher's geometric model. The left-hand panel of each pair shows the mean log fitness of individuals carrying $d$ mutations (eq. 1), and right-hand panel shows the equivalent standard deviation in log fitnesses for individuals carrying $d$ mutations (eq. 2). For all plots, simulations are compared with $k = 1$ (triangles), $k = 2$ (circles) and $k = 3$ (squares). The lines show predictions for the simplest phenotypic model (eqs. 10-11), and the null model (eqs. 3-4 shown as dashed red lines). Each pair of panels shows results from two sets of simulations. (a) the simplest phenotypic models (eqs. 8-9) with $n = 5$ traits, and the two different fitness function (black points: eq. 7; grey points: eq. 6). (b) the fitness function of eq. 6, but with each mutation affecting a distinct trait (black points: $n' = 1$), or a distinct set of 50 traits (grey points: $n' = 50$). (c) the fitness function of eq. 6, with randomly orientated mutations on $n = 5$ traits, whose magnitudes were drawn from a chi-distribution with 0.1 degrees of freedom (black points), or an exponential distribution (grey points). (d) the fitness function of eq. 7, with the effects on each trait drawn from a reflected gamma distribution, with scale parameter 1, and shape parameter $(\sqrt{5}-1)/2 \approx 0.61$ (i.e., a distribution with vanishing mean, unit variance, and a high kurtosis), and results compared for $n = 5$ traits (black points), and $n = 50$ traits (grey points). (e) all details as for panel (d), but with the effects on each trait drawn from a uniform distribution, on the range, $[-0.5, 0.5]$. (f) the fitness function of eq. 6, with $n = 5$ traits, each with a non-zero mean effect of $\beta_i = 0.5$ (black points), or $\beta_i = 0.1$ (grey points). For all simulations, each point shows the mean of $10^6$ replicate individuals, and the strength of selection was scaled, arbitrarily, such that $E(\ln w_1) = -0.01$. When the fitness function of eq. 7 was used, the $\lambda_i$ parameters were generated using the eigenvalues of selection and mutation matrices, which were random Wishart matrices with $n$ degrees of freedom [4,23].

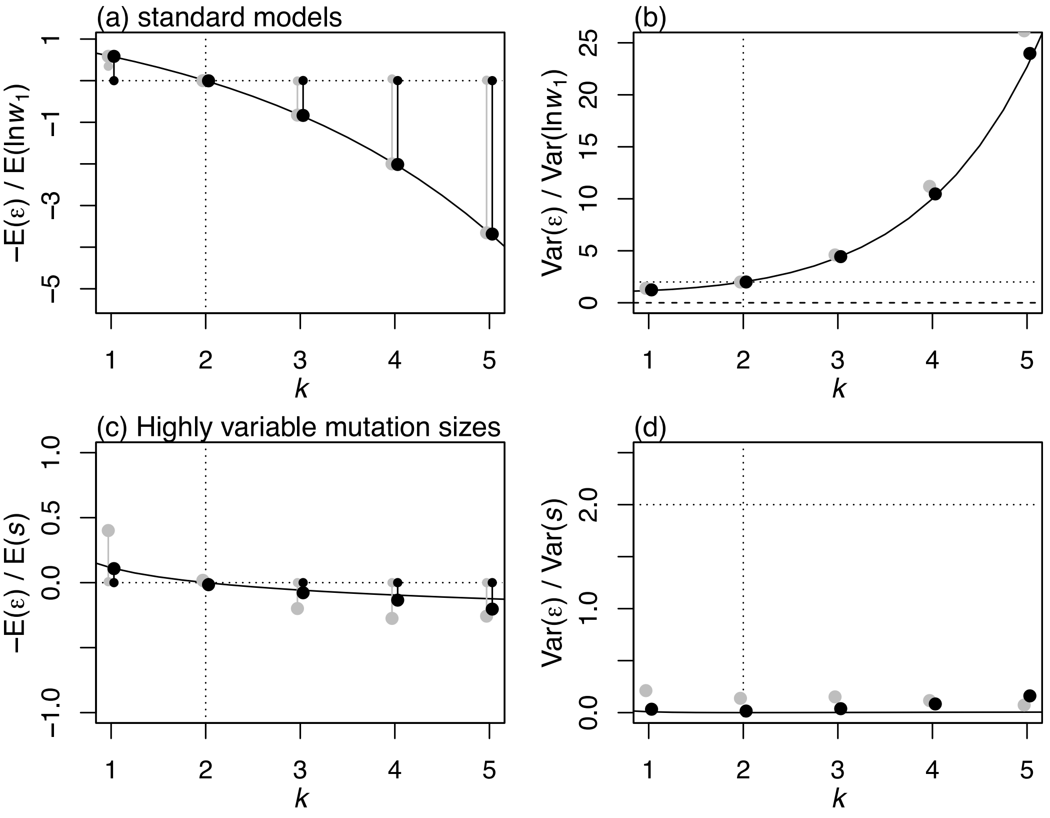


**Figure S2**: Simulations and analytical predictions for the distribution of pairwise epistatic fitness effects (eq. 5), under the additive phenotypic models. Each panel shows the scaled mean or variance in epistasis (eqs. 16-17), as a function of $k$, the curvature of the fitness landscape (eqs. 6-7), and compares predictions (curves) to simulations (points). In panels (a)-(b), mutation effect sizes were normal (eq. 9); curves show eqs. 24-26, and simulations and colours match Figure S1a. In panels (c)-(d), mutation sizes have a highly leptokurtic distribution; curves use eqs. 16, 17, 32 and 33; and simulations and colours match those used in Figure S1c. In panels (a) and (c), larger dots show means, and smaller dots show modal values. To calculate models, we used the half-range mode estimator of Bickel [24] with a bandwidth of 0.95, as implement in the R package *modeest* v. 2.1 [25].

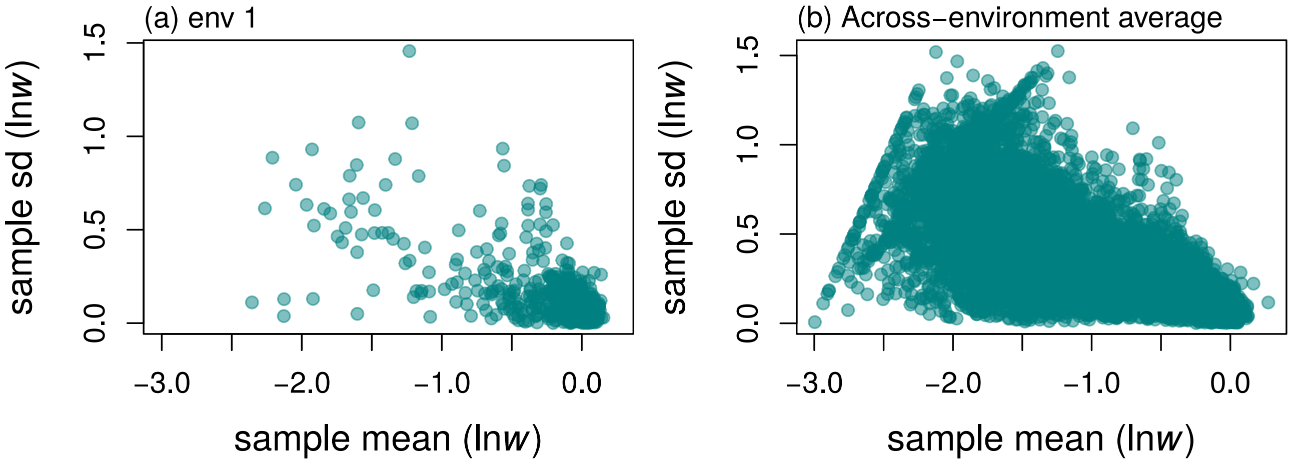


**Figure S3**: The correlation between the mean and standard deviation of replicate measures of mutant fitness for the dataset of Puchta et al. [22]. Results are for all individuals carrying up to $d = 12$ mutations. Panel (a) shows fitness measurements in environment 1, and includes only mutations that were replicated due to multiple hits during the random mutagensis. Panel (b) shows results for all mutations, by treating the 3 environments as replicated measures. The visible lines show the edge effects caused by inability to measure very small fitness values.

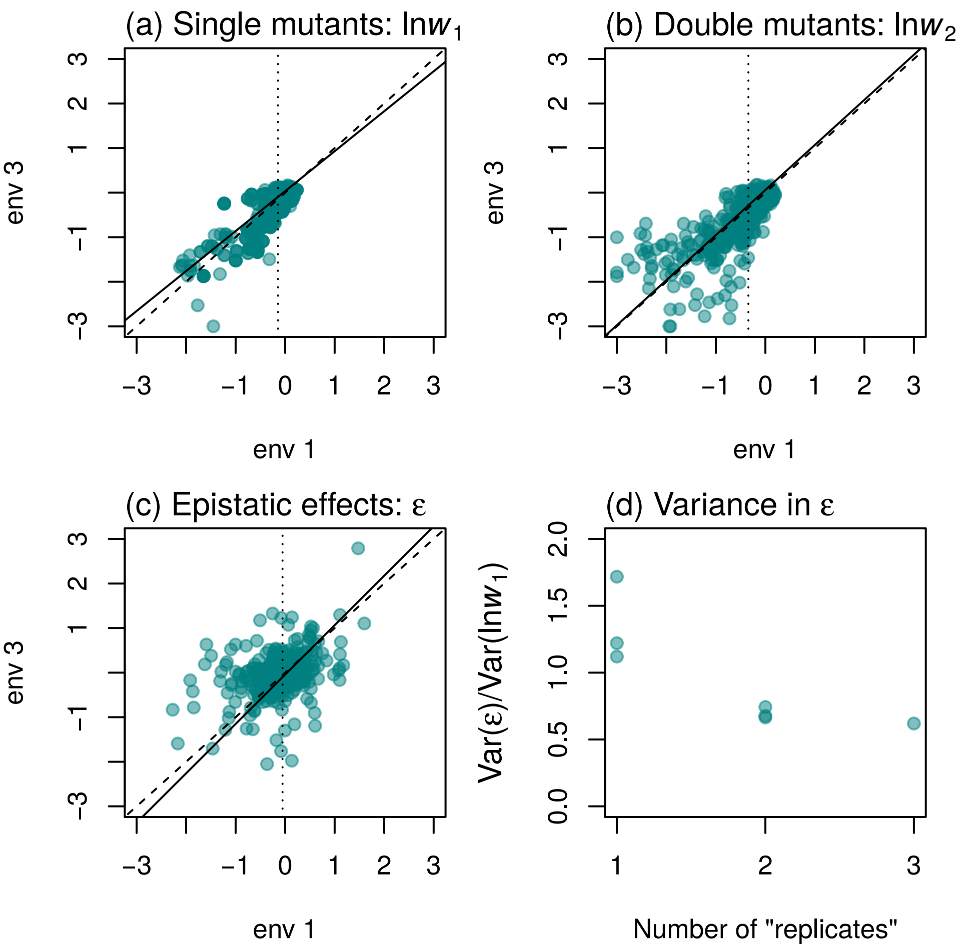


**Figure S4**: *Saccharomyces cerevisiae* snoRNA mutants generated by Puchta et al. [22]. Fitness measurements are shown for the same mutant strains, assayed in two environments, env 1 and env 3 (both comprising glucose 30°C). Results are shown only for Single Nucleotide Polymorphism mutations that were present as both single and double mutants (i.e., discarding all insertions and deletions, and mutations appearing only a singletons). Panel (a) shows the single mutants; panel (b) the double mutants, and panel (c) shows the corresponding epistatic effects (eq. 5). In each case, the best-fit Standardized Major Axis regression (solid line) is compared to the 1:1 slope (dashed line). Panel (d) shows the scaled variance in epistatic effects (eq. 17), when the log fitness values were either measured in a single environment, or averaged over 2 or 3 environments. Increasing the level of replication decreases the inferred variance in epistatic effects.

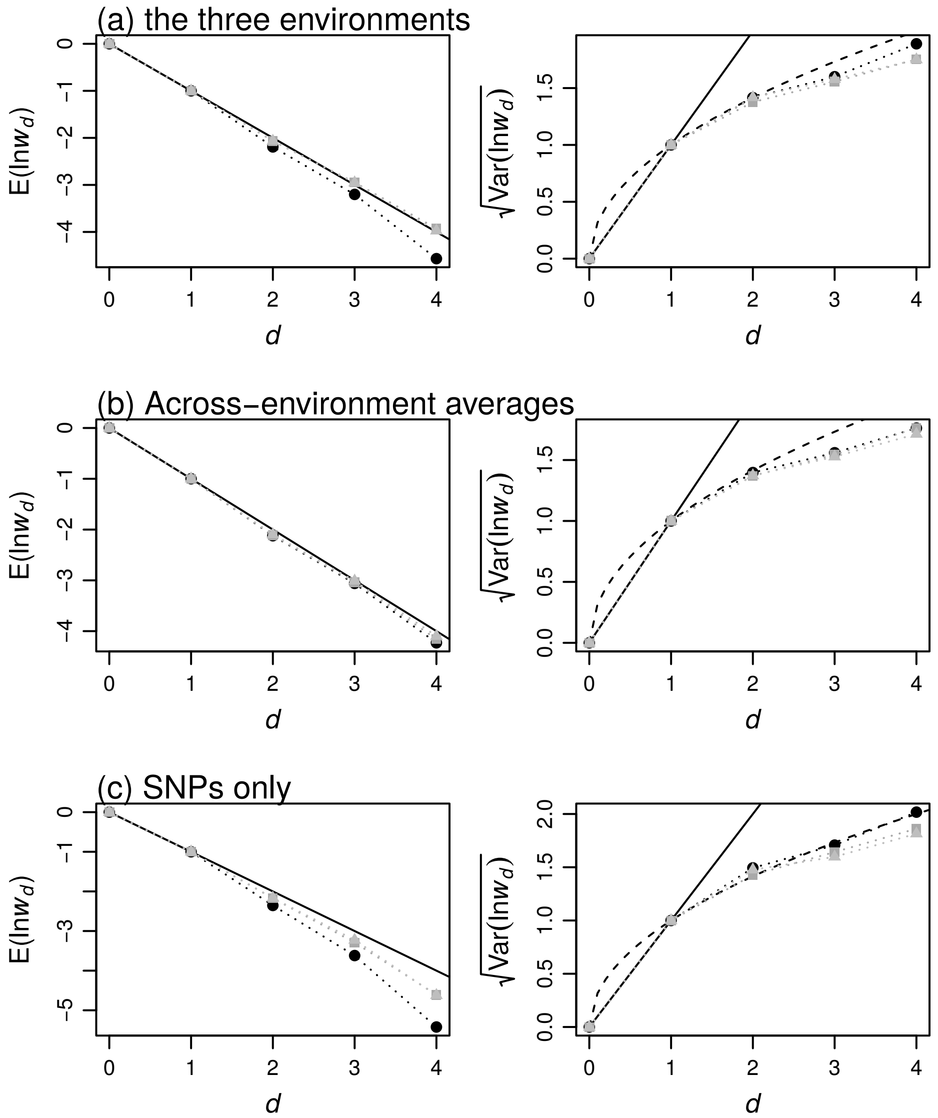


**Figure S5**: *Saccharomyces cerevisiae* snoRNA mutants generated by Puchta et al. [22], and assayed in competition experiments in three environments (env. 1 and 3 in glucose at 30°C, and env. 2 in glucose at 37°C). All plots show the mean and standard deviation in the log fitnesses of individuals carrying $d$ mutations, as in Figure 2. Panel (a) shows results for the three environments separately (env. 1: black circles, env. 2: dark grey squares, and env. 3: lighter grey triangles). Panel (b) shows results when log fitness measurements were averaged across environments: (env. 1 and 3: black points, env. 1 and 2: dark grey squares, and all three environments: lighter grey triangles). Panel (c) is identical to panel (a), but shows only Single Nucleotide Polymorphism mutations (i.e., discarding small insertions and deletions).

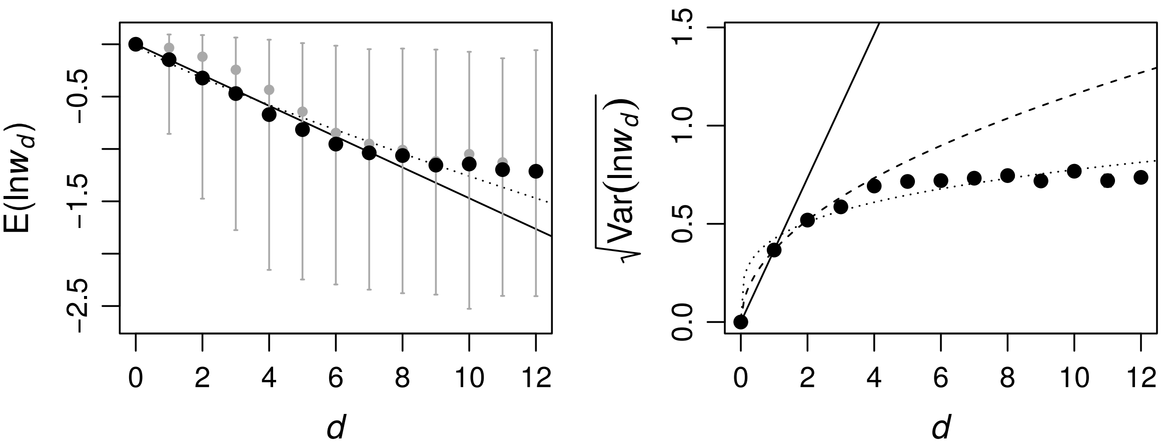


**Figure S6**: *Saccharomyces cerevisiae* snoRNA mutants generated by Puchta et al. [22]. Plots are identical to Figure 2c-d, but show results for individuals carrying up to $d = 12$ mutations. Edge effects, caused by the inability to measure fitness accurately below a certain value, have a visible effect after the first few mutations. This explains why our main results were truncated at $d = 4$.
